# Computational response modeling reveals context dependent Akt activity in luminal breast cancer cells

**DOI:** 10.1101/2020.10.22.349647

**Authors:** Cemal Erdem, Adrian V. Lee, D. Lansing Taylor, Timothy R. Lezon

## Abstract

Aberrant signaling through insulin (Ins) and insulin-like growth factor I (IGF1) receptors contributes to the risk and advancement of many cancer types by activating cell survival cascades. Mechanistic computational modeling of such pathways provides insights into each component’s role in the cell response. In previous computational models, the two receptors were treated as indistinguishable, missing the opportunity to delineate their distinct roles in cancer progression. Here, a dual receptor (IGF1R & InsR) computational model elucidated new experimental hypotheses on how differential early responses emerge. Complementary to our previous findings, the model suggested that the regulation of insulin receptor substrate (IRS) is critical in inducing differential MAPK and Akt activation. As predicted, perturbing ribosomal protein S6 kinase (RPS6K) kinase activity led to an increased Akt activation with insulin stimulation compared to IGF1 stimulation. Being able to discern differential downstream signaling, we can explore improved anti-IGF1R cancer therapies by eliminating emergence of compensation mechanisms, without disrupting InsR signaling.

**Implications:** The study shows, both experimentally and through computational models, that IGF1 and insulin receptor signaling pathways respond differently to RPS6K inhibition.

## INTRODUCTION

Insulin and type I insulin-like growth factor (IGF1) are closely related hormones that are critical to development and metabolism (1–4). Their receptors, InsR and IGF1R, are structurally and functionally similar, showing 60% overall amino acid sequence similarity and 84% identity at the kinase domain (5,6). The signaling pathways of both receptors have been linked to cancer, where both can activate proliferation and survival cascades. Increased insulin and IGF1 levels have been shown to correlate with increased risk of several cancer types (7–9). IGF1R content in breast cancer (BRCA) tumors is 14 times higher than in normal tissue, and inhibiting IGF1R has been shown to block tumor growth in cell lines and model organisms (10–15).

While no recurrent cancer-specific mutations of IGF1R or its ligands have been described to date, studies have provided evidence for a link between this signaling pathway and the risk of developing cancer. IGF1R signaling leads to both proliferative and anti-apoptotic signaling by employing Ras/MAPK and PI3K/Akt cascades. Clinical trials of IGF1R targeting received some positive responses; however, compensation mechanisms emerge and decrease the efficacy of such drugs (16–18). Insulin stimulates cell growth, differentiation, and promotes synthesis while inhibiting lysis of macromolecules (4). Insulin malfunctioning results in dysregulation of these processes and causes elevated glucose and lipid levels. Insulin resistance has been associated with both type II diabetes and obesity, where increased insulin levels is shown to correlate with increased risk of several cancer types (4,7,8,19).

The insulin and IGF1 receptors are heterotetramers, or rather a dimer of heterodimers, with two α and two β subunits. Beta subunits contain the intracellular kinase domains. The α-subunits span the extracellular ligand binding domains. IGF1R and InsR can also form hybrid receptors, with one α-β pair from each. These hybrid receptors show differential affinity for the three ligands (20). Two recent studies suggested that the extracellular domains (ECD) of the apo-receptor forms exert a physical force to keep the intracellular kinase domains apart from each other, enough to prevent auto-phosphorylation (21,22). Ligand binding then induces a conformational change that lets the transmembrane and kinase domains to interact and auto-phosphorylate. The changes upon ligand binding are studied by Houde and Demarest in (23). Kiselyov et al. used modeling approaches to re-capture available ligand binding dynamics of insulin in (24). This study only considered the ligand-receptor binding events, ignoring the fact that downstream elements of the transduced signal also affect available receptors on the cell surface.

An understanding of the differences between the highly similar InsR and IGF1R signaling is important for therapeutic development and clinical trial design. In a previous study (25), we constructed statistical models of insulin and IGF1 signaling from a large proteomics data set. Our models revealed, and experiments confirmed, cell-level differences in signaling through the two pathways. Specifically, acetyl-CoA carboxylase (ACC) or E-Cadherin knock-down increases MAPK or Akt phosphorylation, respectively, in IGF1 stimulated cells over Ins stimulated cells (25). Although subsequent work showed that loss of E-Cadherin increases sensitivity of breast cancer cells to IGF1R/InsR targeted therapy by hyperactivating the IGF1R signaling pathway (26), the precise mechanisms through which ACC and E-cadherin influence signaling remains hidden.

Here we construct a mechanistic model of IGF1R/InsR signaling to identify downstream differences in signaling through the two receptors. Most existing models treated IGF1R and InsR signaling identically (27–30), while some, although they modeled them individually, did not focus on studying them (31). In contrast, our model retains each receptor’s unique identity and recovers differences in signaling through IGF1 and Ins. We show through systematic parameter scanning that perturbing ribosomal protein S6 kinase (RPS6K) activity should increase Akt activation with insulin stimulation compared to IGF1 stimulation, and we experimentally confirm this prediction. Our work demonstrates that modulating targets downstream of IGF1R and InsR may provide an alternative to specifically modulating IGF1R, potentially allowing targeted IGF1R therapies that do not disrupt insulin signaling and glucose metabolism.

## METHODS

### Initial dataset and computational model parameter estimation

The initial dataset utilized here was defined in our previous work (25). In short, reverse phase protein array (RPPA) is a high-throughput technique for quantifying levels of total and phosphorylated proteins (32,33). The model parameters were estimated using RPPA expression levels of four phospho-proteins: pReceptor (both IGF1R and InsR), pAkt, pRPS6K, and pMAPK. Data from three early time points (5, 10, and 30 min) were used to estimate the parameters and to calculate the fitting error.

In this work, all ODEs corresponding to the state variables were obtained from BioNetGen (34), and the rest of the simulations and analyses were done in MATLAB (The MathWorks, Inc., version R2015a). The parameter estimation was done using Markov chain Monte Carlo (MCMC) sampling, at different temperatures to search the parameter space both locally and globally (35). High temperature chains scan the parameter space more globally, and the probability of accepting an unfavorable move depends on the temperature. The swaps among different chains help avoid getting stuck in local minima. The approach samples the Bayesian posterior distribution of each parameter, with uniform priors (36,37). The estimation procedure outputs parameter ensembles for each chain. The minimum fitting error parameter set was defined as the “best-fit” and was used for all subsequent analyses.

### *In-silico* ensemble of cells

Using the value-ranges set for the initial protein count parameters, an ensemble of 10,000 parameter sets were generated using Latin hypercube sampling. By only changing the values of total protein numbers and keeping estimated rate parameters constant, different cell conditions were captured (i.e. a virtual cell population).

### Parameter perturbation scanning

Once the parameters were determined, simulations were run to analyze the response of the system. Each parameter was perturbed individually and for every different value of each parameter, one simulation was run. The predicted levels of pMAPK and pAkt in the perturbed system were compared to the levels in the un-perturbed model output, simulating experimental knock-down (up-regulation) of proteins or reaction rates. Based on the results of the simulations here, perturbations that resulted in differential responses under IGF1 and insulin stimulation conditions were selected for further experimental exploration.

### Cell culture and immunoblotting

MCF7 (ATCC) cells were cultured in DMEM (ThermoFisher) with 10% FBS, plated on six well plates at 400000 cells/well density. The cells, rested overnight, were serum starved for 16-24 hours. Then, the cells were treated with DMSO control or ribosomal protein S6 kinase inhibitor LY2584702 (500 nM, Selleckchem) for three hours. Next, the cells were stimulated with control, IGF1 (10 nM), or insulin (10 nM) for 10 and 30 min. The cells were harvested, and protein concentration was quantified by BCA. Samples were collected using RIPA buffer (50mM Tris pH 7.4, 150mM NaCl, 1mM EDTA, 0.5% Nonidet P-40, 0.5% NaDeoxycholate, 0.1% SDS) with 1x HALT protease & phosphatase inhibitor cocktail (ThermoFisher). The immunoblotting was done using 12% acrylamide gels and PVDF membrane transfer (Millipore #IPFL00010, 0.45µm). Membranes were blocked in Odyssey PBS Blocking Buffer (LiCor), and incubated in primary antibodies overnight: Akt S473 (Cell Signaling #4060; 1:1000), total Akt (Cell Signaling #2920; 1:1000), phospho-S6 S235/236 (Cell Signaling #4858, 1:1000), total S6 (Cell Signaling #2217, 1:1000), and β-actin (Sigma #A5441; 1:5000). Membranes were incubated in LiCor secondary antibodies for 1 hour (anti-rabbit 800CW, LiCor #926-32211 or anti-mouse 680LT, LiCor #925-68020; 1:10000) at room temperature. The imaging was done at LiCor Odyssey Infrared Imager, where blots were quantified using LiCoR Image Studio Lite v5.2 software.

T47D (ATCC) and ZR75-1 (ATCC) cells were cultured in RPMI-1640 (HyClone, GE) with 10% FBS, 1% glutamine, and 1% Penicillin-Streptomycin, plated on six well plates at 500000 cells/well density. The cells, rested overnight, were serum starved for 16-24 hours. Then, the cells were treated with DMSO control or ribosomal protein S6 kinase inhibitor LY2584702 (1 μM) overnight. Next, the cells were stimulated with control, IGF1 (10 nM), or insulin (10 nM) for 10 and 30 min. The cells were harvested, and protein concentration was quantified by Bradford absorbance assay. Samples were collected using HEPES buffer (1% Triton X-100, 10% Glycerol, 5mM MgCl2, 25mM NaF, 1mM EGTA, 10mM NaCl) with 1x HALT protease & phosphatase inhibitor cocktail (ThermoFisher). The immunoblotting was done using 12% acrylamide gels (ThermoFisher #XP00125BOX) and PVDF membrane transfer (ThermoFisher #LC2002, 0.2 µm). Membranes were blocked in 5% milk in 1X TBST solution (TBST: Tris Buffered Saline (Sigma # T6664) with 0.1% Tween20), and incubated in primary antibodies overnight: Akt S473 (Cell Signaling #4060; 1:1000), total Akt (Cell Signaling #2920; 1:1000), phospho-S6 S235/236 (Cell Signaling #4858, 1:1000), total S6 (Cell Signaling #2217, 1:1000), and β-actin (Sigma #A5441; 1:5000). Membranes were incubated in HRP secondary antibodies for 45 min at room temperature (anti-rabbit, Jackson #111-035-003 or anti-mouse, Jackson #115-035-003; 1:8000). The imaging was done on a Philipps L4000 Imager using ECL substrates (BioRad #170-5060). The blots were quantified using LiCor Image Studio Lite v5.2 software.

All three cell lines are parental cells obtained from ATCC and internally mycoplasma tested regularly or when suspected of any contamination. No such observation during this study.

## RESULTS

### Computational model

We developed our model (Fig. 1A) from previous work in which only IGF1R signaling (27,28) or InsR signaling (29,30) was modeled. Ligand binding to IGF1 or insulin receptors promotes receptor intracellular domains to auto-phosphorylate, leading to IRS and SOS binding and their activation (27–29,38,39). Phosphorylated IRS can also activate SOS in addition to PI3K (29,40,41). SOS activation leads to activation/phosphorylation of Ras, Raf, MEK, and MAPK (ERK) (10,27). PI3K activation causes PDK1, Akt, mTOR, and RPS6K activation. There are numerous negative feedback loops and crosstalk within the system (Fig. 1A) (28,42).

**Figure 1.**
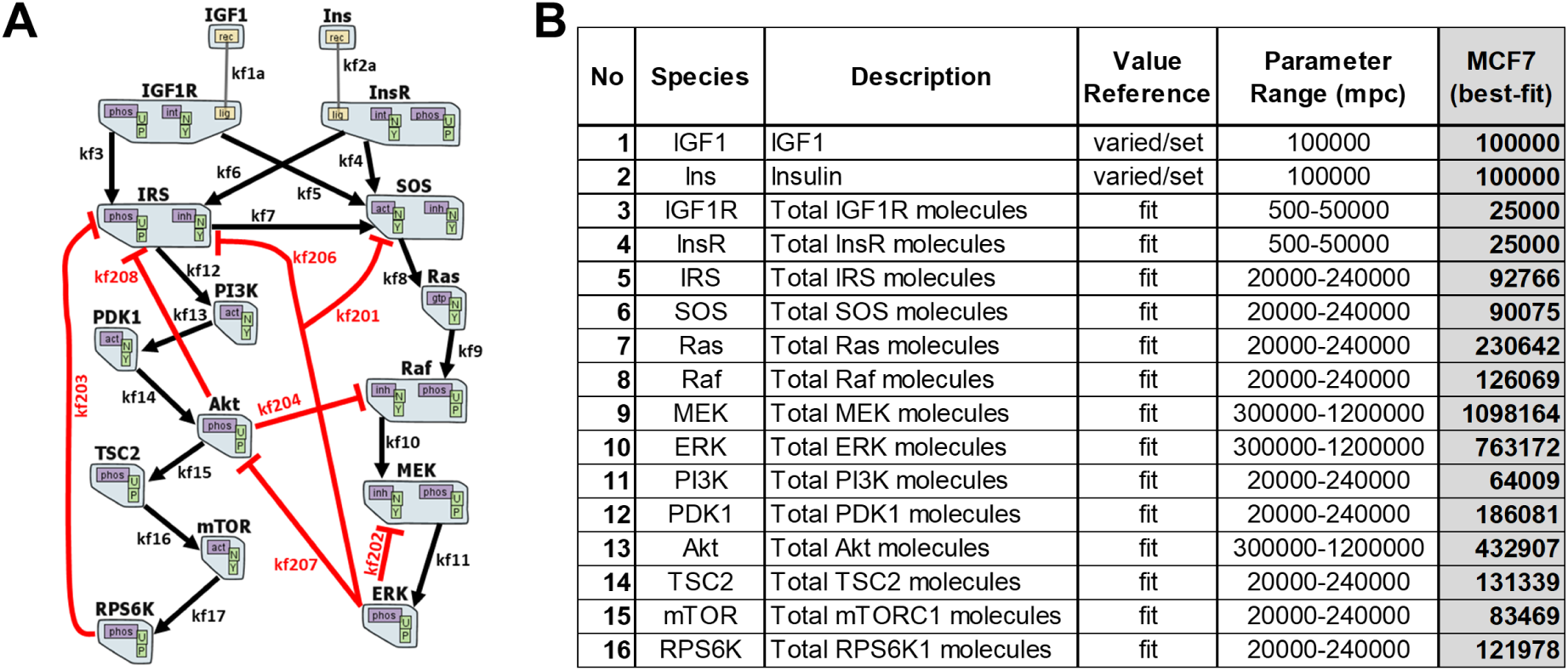
The computational mechanistic model representation. (A) The topology of dual IGF1R/InsR signaling network is illustrated. The model includes 16 proteins. Black arrows represent activation and red lines indicate inhibition of the corresponding active molecule. The graph is adapted from RuleBender software. (B) The initial molecule numbers of the species in the model. The parameter value ranges and the “best-fit” values are reported. mpc: molecules per cell.

The constructed model has 14 proteins and two ligands, and 66 parameters, of which 16 are the total protein counts (Fig. 1A and Table S1). The protein counts, and 34 of the rate parameters, are common between IGF1 and insulin models, where each model has eight specific parameters. See BNGL model in Supplemental Text 1, and corresponding state variables and ODEs in Supplemental Text 2. The mechanistic model in this work was constructed using rule-based modeling with BioNetGen and RuleBender software (34,43–45).

### The computational training performance of the model

The experimental dataset used for model training contains proteomic (RPPA) data of phosphorylated and total protein levels, at six different discrete time points (0-48 hours). Actually, most of the activation (or phosphorylation) events in the downstream cascades occur within minutes post-treatment. By studying signaling in the earlier time points, we hope to unravel some *early response differences* of the breast cancer cells to single IGF1 or insulin stimulation.

The parameter estimation was performed using Markov chain Monte Carlo (MCMC) sampling and parallel tempering (35). The procedure yields ensembles of parameter sets and corresponding posterior distributions (Figs. S1 and S2). The resulting parameter set (Table S1) of the network model was estimated and simulation results conveyed qualitative and quantitative agreement to experimental data for both IGF1R and InsR signaling (Fig. 2A). The dashed lines represent the performance of the best-fit parameter set model on training data. In addition to the best-fit models of IGF1 and insulin stimulation, the ODE models were simulated for an ensemble of parameter sets (area plots in Fig. 2A). This corresponds to screening a virtual population of cells, where each parameter set has different initial conditions set for the simulation. Within the specified range of parameters, the computational model recaptures experimental data, indicating a decent set of rate parameters have been estimated.

**Figure 2.**
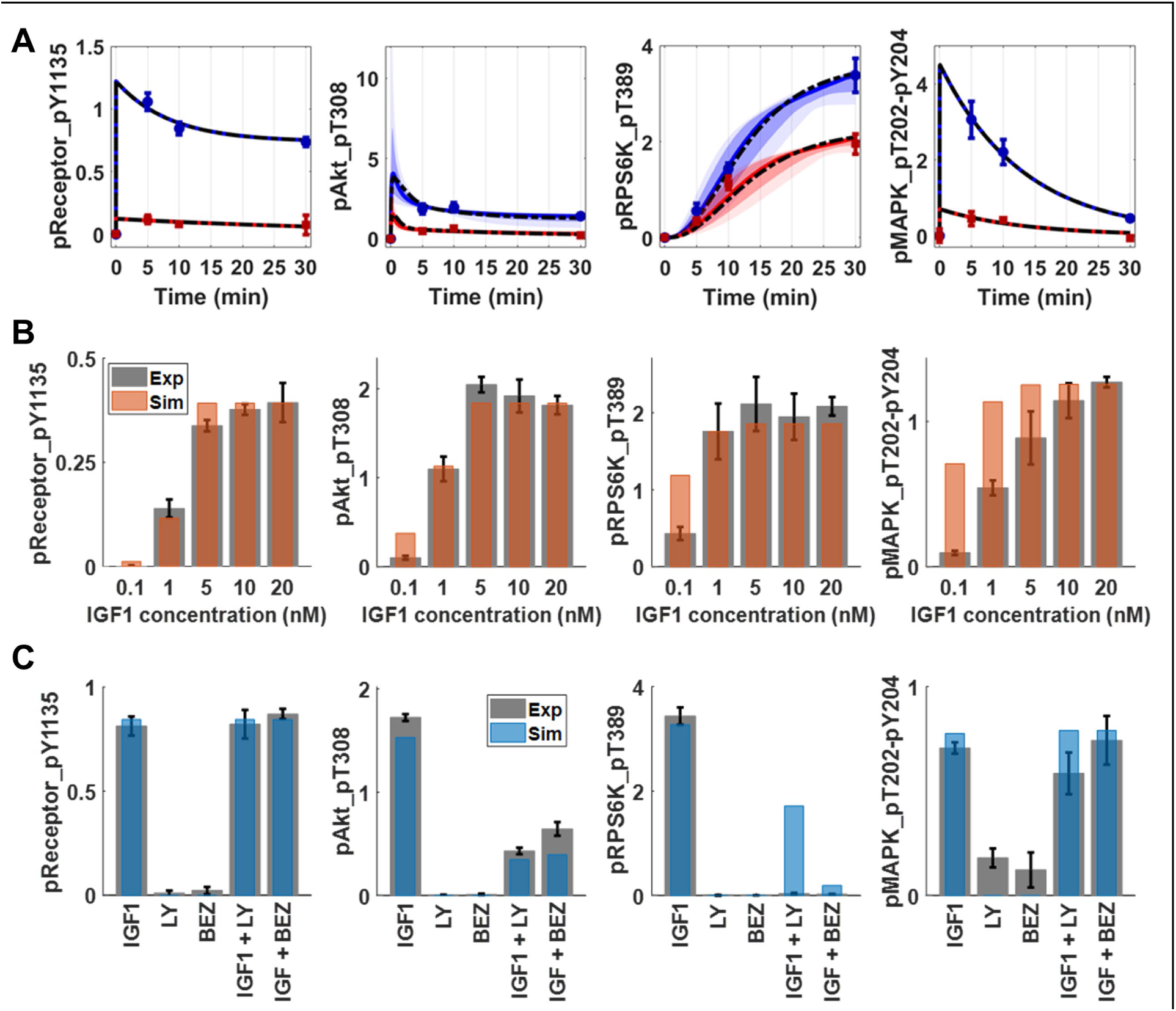
The mechanistic model training and test performance on multiple omics datasets show qualitative and quantitative agreement. (A) The time course trajectories of IGF1 (blue) and insulin (red) stimulations. The ODE model was simulated 10000 times with an ensemble of parameter sets, with different total protein numbers. The initial amounts of proteins were sampled using Latin Hypercube Sampling. The plots show 5%-95% (light) and 15%-85% (dark) confidence intervals for IGF1 (blue) and insulin (red) models. The circles with error bars are the corresponding RPPA data points. The dashed black lines are the trajectories for “best-fit” parameter sets. (B) The computational model (orange bars) recapitulates experimental IGF1 dose response data (gray bars). The error bars represent the standard error of the mean from three independent biological replicates. The y-axis represents scaled protein numbers. (C) The computational model (blue bars) recapitulates PI3K inhibition data (gray bars). The columns of x-axis correspond to: IGF1: IGF1 (10 nM) stimulation, LY: first inhibitor only, BEZ: second inhibitor only, IGF1+LY: first inhibitor and IGF1 (10 nM), and IGF1+BEZ: second inhibitor with IGF1 (10 nM) stimulation. The receptor and MAPK phosphorylation are not affected by PI3K or mTOR inhibition whereas Akt and S6 kinase phosphorylation are decreased. The inhibitions are simulated in the computational models as a 90% reduction in the corresponding rate constant(s). The error bars represent the standard error of the mean from three independent replicates. The y-axis represents scaled protein numbers.

Additionally, a sensitivity analysis of the model was carried out and details are described in Supplemental Text 3. The cascade-specific parameters affected the corresponding readout the most, and the parameters of receptor kinetics induced changes in receptor phosphorylation levels (Fig. S3-6).

There are other mechanistic models in the literature (28,31,46), spanning different proteins and interactions (edges) among the protein species in our model. We specifically tested individually adding the interactions depicted in (28) into our model. Parameter scanning these model variants did not yield an improvement in data fitting and thus the candidate interactions are discarded from the final model topology.

### IGF1 dose response and PI3K inhibition in MCF7 cells

We tested the model performance using two independent datasets: IGF1 dose response (Fig. 2B, Table S2) and PI3K/mTOR inhibition (Fig. 2C, Table S3). The simulation results for IGF1 dose response were obtained by only changing the level of IGF1 input into the system. The results were within the range of experimental error and showed qualitatively good agreement with the data.

A second tier of performance test was done using PI3K and mTOR inhibition data. One specific PI3K inhibitor, LY294002 (LY), and one dual inhibitor of PI3K and mTOR, BEZ235 (BEZ), were administered alone or in combination with IGF1 (10 nM) (Table S3). To simulate the activity of the PI3K inhibitor LY294002, the rate parameter controlling PI3K-mediated activation of PDK1 was decreased by 90%. To simulate the activity of the PI3K/mTOR inhibitor BEZ235, this rate constant and the constant that controls S6K activation through mTOR were both decreased by 90%. Without any ligand input, neither inhibitor affected any downstream signaling (Fig. 2C gray bars-experimental data, blue bars-simulations). Addition of IGF1 into the inhibitor-treated system produced decreased phosphorylation of Akt compared to treatment with IGF1 alone. Although ribosomal protein S6 kinase phosphorylation was diminished in both inhibitor experiments, the computational model predicted a non-zero activation for IGF1 stimulated cells with PI3K inhibitor LY. Even with 90% inhibition of PI3K, minimal mTOR activation occurred and led to phosphorylation of S6K. The experimental observation might be a result of the fact that the kinase inhibitors are dirtier than the simple perturbations simulated here (47). The model recapitulated full deregulation of S6K phosphorylation in the case of another inhibitor (BEZ) with multiple targets. Overall, the model showed qualitative agreement with the experimental results, and captured expected behavior of the perturbed systems, with minimal parameter perturbations.

### Simulation results reveal details of known IGF1-strong responses

We next scanned parameter values to search for specific perturbations that capture the effects that we observed in our earlier knock-down experiments of ACC (acetyl-coA carboxylase) and E-Cadherin (25). The first analysis was carried out to determine clues for ACC action on MAPK phosphorylation. Previously, it was shown that the ACC knock-down causes an increase in MAPK activation, with a larger change induced by IGF1 than by insulin. The parameter perturbation scan results showed that the upregulation of the rate of SOS activation by IRS protein (rate parameter k7) should be explored further to pinpoint the mechanism of action of ACC on MAPK (Fig. 3A top panel). It is of note that the differential response was captured by “up-regulation” of a rate, rather than a knock-down.

**Figure 3.**
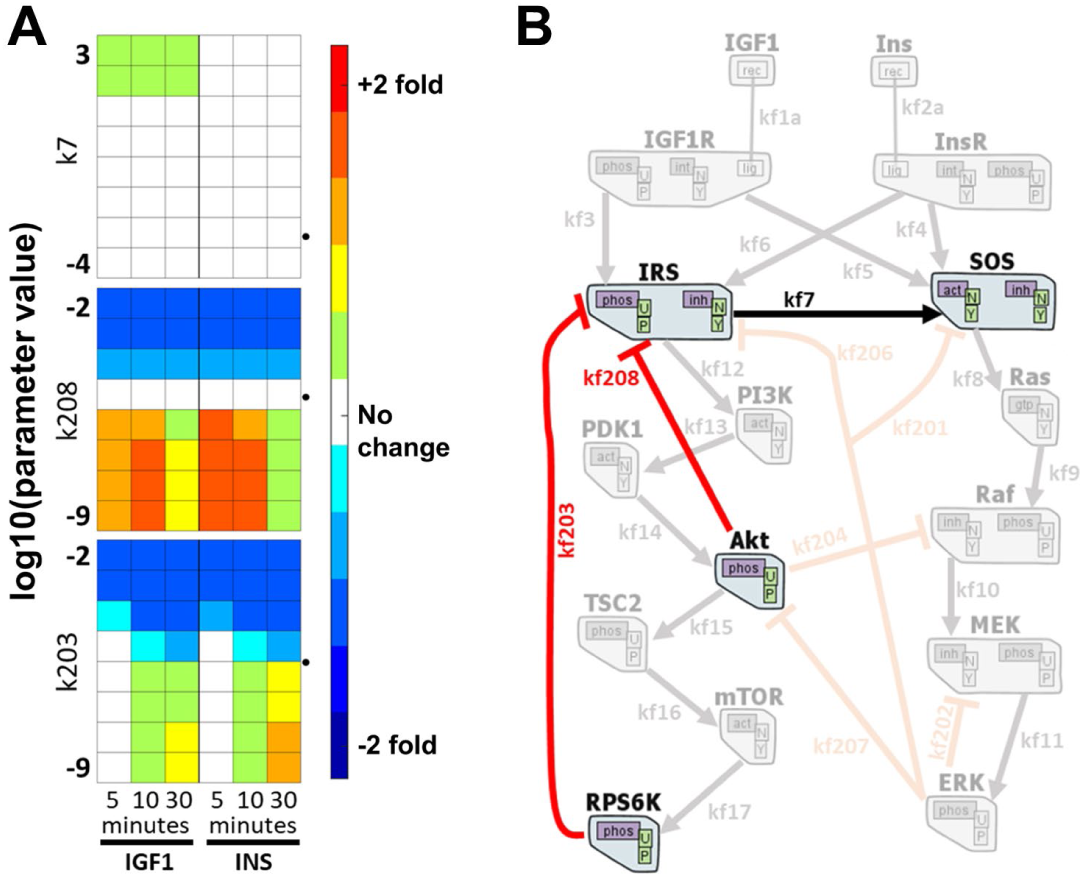
*In silico* perturbation of rate constants reveal differential effects on Akt and MAPK phosphorylation. The model output of pMAPK and pAkt levels are determined by setting and changing the value of each parameter individually, from zero to infinity. Each box above is for a parameter, and each column is for one time point response of 5, 10, and 30min. Left three columns are from IGF1 stimulation results and the right three columns are from insulin stimulated simulations. Rows represent the log10 of set parameter value. The colors represent the log2 fold change from the un-perturbed model output. Red and blue respectively indicate up and down regulation of the specified phosphorylation. The dotted arrow heads indicate the unperturbed model parameter values in log10. Rest of the parameter perturbation scanning results are shown in Fig. S7. (B) The respective edges from (A) are depicted together to emphasize that all perturbations leading to differential regulation are concentrated on IRS proteins.

Second, changing the value of each parameter individually and analyzing the resulting changes in Akt phosphorylation levels. We found that disrupting the negative feedback of Akt on the upstream adaptor protein IRS had the same effect on Akt activation as did E-Cadherin knock-down. Turning off rate parameter kf208, corresponding to the feedback of Akt on IRS, caused differential up-regulation of Akt activation (Fig. 3A middle panel). A larger increase at 30min with IGF1 stimulation was verified experimentally, recapitulating our earlier observations in E-Cadherin knocked-down cells (25).

### Mechanistic modeling predictions reveal *novel* insulin-strong responses

Differential regulation of Akt and MAPK phosphorylation were further explored based on the results of parameter perturbation scanning. One of the predictions with a differential response from IGF1 and insulin is the knock-down of ribosomal protein S6 kinase. We predicted that upon inhibiting S6 kinase, the insulin stimulated cells would have increased Akt phosphorylation at 30 min, and that the magnitude of the increase would be larger than that in the IGF1 stimulated cells (Fig. 3A bottom panel). The rate parameter “kf203” controls negative feedback from S6 kinase on IRS. All three computational predictions pointed out that regulation of IRS is critical for differential downstream regulation of IGF1 and insulin receptor cascades (Fig. 3B).

#### Ribosomal protein S6 kinase (RPS6K) inhibition in luminal BRCA cells

We experimentally validated the last prediction by chemically inhibiting ribosomal protein S6 kinase in breast cancer cells, and then treating with IGF1 and insulin, as described in *Methods* The experiments were performed in MCF7, T47D and ZR75-1 cell lines, all of which are luminal and hormone receptor positive subtype. The decrease in ribosomal protein S6 phosphorylation (by RPS6K) levels was greatest in MCF7 cells (Fig. 4), but in all three cell lines, the level of Akt phosphorylation (S473) increased in insulin stimulated drug treated cells, compared to IGF1 stimulated cells. This result follows the computational prediction of the mechanistic model.

**Figure 4.**
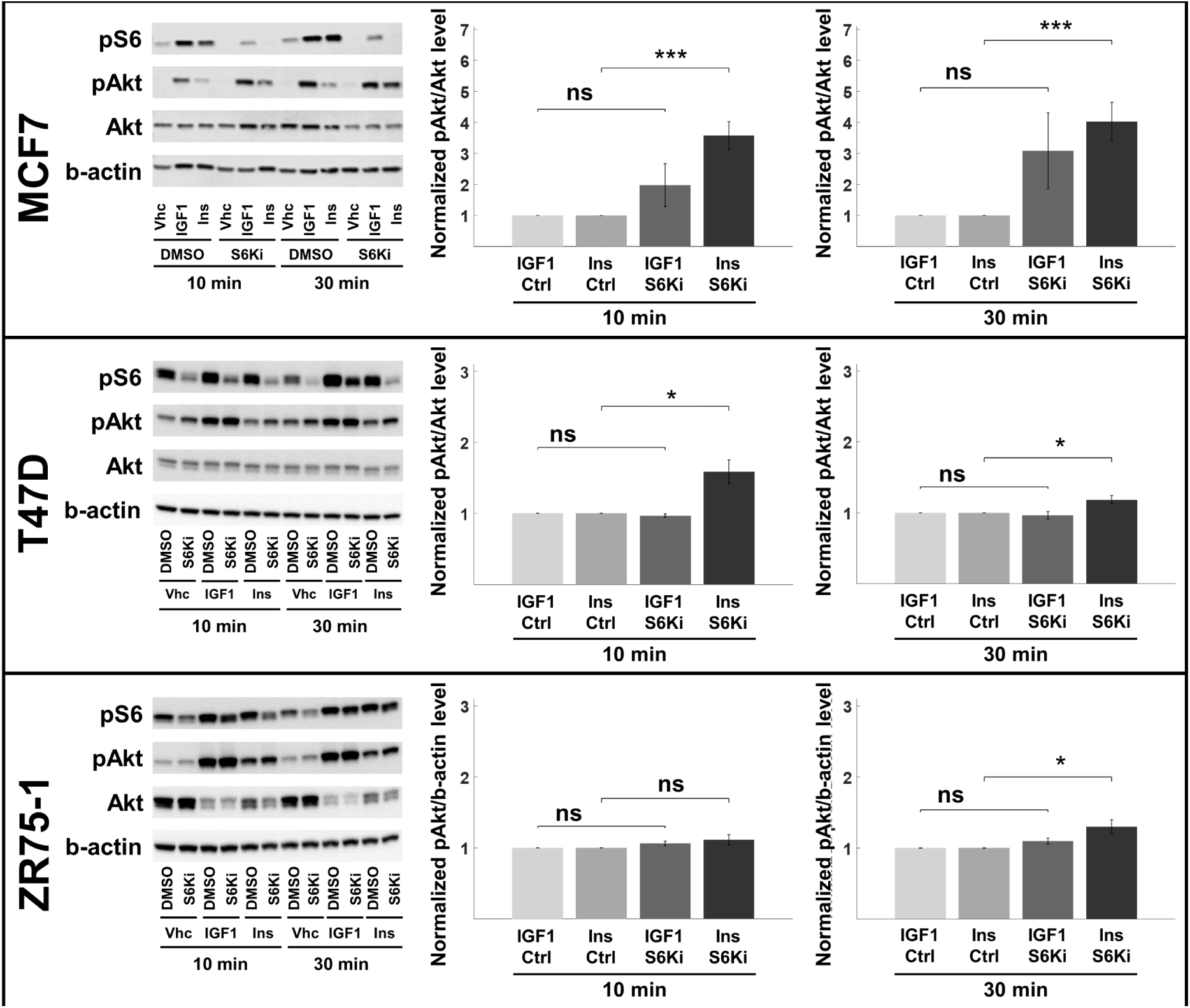
Ribosomal S6 kinase inhibition up-regulates Akt phosphorylation. The RPS6K was inhibited in MCF7, T47D, and ZR75-1 cells. The pS6 levels are used as the proxy for S6K inhibition efficiency. The total Akt and b-actin levels is used to normalize pAkt and pS6 levels, respectively. The perturbagen, ligand, and time point for each sample are listed below the blot images (rest of the replicates are shown in Fig. S8), and the blots are quantified. The values are reported as normalized to the corresponding no-inhibition control (ctrl). pAkt levels represent the response of the cells to the perturbation. All MCF7, T47D, and ZR75-1 cell lines showed higher up-regulation of pAkt in insulin stimulated cells at 30 min. The results are compared using unpaired, one-tailed two-sample t-test, and P<0.05 (^*^), P<0.01 (^**^), P<0.005 (^***^), nonsignificant (ns). Results shown are mean ± s.e.m. of four independent replicates. S6 phosphorylation quantifications at 10 and 30 min are shown in Fig. S9.

## DISCUSSION

In our earlier work, we showed that acetyl-CoA carboxylase knock-down increases MAPK phosphorylation while E-cadherin knock-down promotes higher Akt activation in IGF1 stimulated cells (25). These were novel findings that show how insulin and IGF1 downstream signaling cascades differ in cancer cells. In this work, we used a detailed mechanistic model to generate new experimental hypotheses on how ACC and E-Cadherin knock-downs result in distinct responses upon IGF1 and insulin administration. The results here suggest that the stronger response of MAPK to IGF1 than to insulin depends critically on SOS activation through IRS. For the E-Cadherin knock-down data, the negative feedback from Akt to IRS is important to obtain the differential Akt activation. Mechanisms of action of both ACC and E-Cadherin are hypothesized to focus on the regulation of IRS, which is an adapter protein and one of the bottlenecks of signaling activation (48,49). Recent structural analysis of the two RTKs also suggests a differential binding of IRS proteins (39).

*In the current work, the feedback from ribosomal protein S6 kinase (p70S6K) on IRS is predicted to differentially affect Akt activation under IGF1 and insulin stimulated cells*. The experimental validation of the *in-silico* prediction in three different cell lines shows that there indeed is a difference in the regulation of Akt activity in response to different stimuli, with a greater change induced by insulin rather than by IGF1. This result indicates a tighter regulation of IRS by RPS6K in insulin stimulated cells, such that revoking the negative feedback causes a larger up-regulation of Akt phosphorylation in hormone receptor positive luminal breast cancer cells. In addition, relieving the negative feedback from S6K on IRS was previously shown to sensitize colorectal cancer cells to EGFR inhibition (28). Similarly, our collaborators studied IGF1R/InsR pathway sensitization in the absence of E-Cadherin (26). These results all convey the importance of regulation on IRS and how it activates downstream cascades in different cellular contexts.

We studied how IGF1 and Ins activate their cognate receptors. The receptors actually have different isoforms, IGF1R / IGF2R and InsR-A / InsR-B (20). IGF1R binds to both IGF1 and IGF2 (insulin-like growth factor 2). The latter ligand is mostly fetal and not studied in this work. IGF2R, the IGF1R homolog, has no kinase domain and considered to sequester IGF2 primarily (20). One isoform of insulin receptors, the InsR-A, is functional in fetal tissues and cancer cells, whereas InsR-B isoform is expressed in adults. InsR-A has similar affinity for IGF2 and insulin (6,11). There are even *heterodimers* of IGF1R/InsR (20), which can ideally bind any of the ligands. In this work, we only focused to reveal how IGF1 and Ins stimulations exert different responses in breast cancer cells and not focused on the isoform specificity.

The study of system biology encompasses employment of tools and techniques to extract information from large datasets. Within the quantitative systems pharmacologic (QSP) framework undertaken here, the mechanistic computational model is supplied with experimental and observational data and is iteratively refined (50–52). The role of the models was then to simulate different stimulation conditions *in silico* and analyze the response with any possible off-target effects. In doing so, we can start stratifying patients to suitable personalized medicine treatments after recognizing and distinguishing that the IGF1R and InsR systems have different dynamics and novel signaling components.

## ACKNOWLEDGEMENTS

CE gratefully acknowledges Dr. Alison Nagle & Beth Knapick from Lee lab, and Laura Vollmer & Celeste Reese of UPDDI for wet-lab training. This work was funded by P30 CA047904 and U01CA204826 (DLT), and NIH 1UL1TR001857-01 (TRL).

## REFERENCES

1. Liu JL, LeRoith D. Insulin-like growth factor I is essential for postnatal growth in response to growth hormone. Endocrinology. 1999;140:5178–84.

2. Braun S, Bitton-Worms K, LeRoith D. The Link between the Metabolic Syndrome and Cancer. Int J Biol Sci. 2011;7:1003–15.

3. Kooijman R. Regulation of apoptosis by insulin-like growth factor (IGF)-I. Cytokine Growth Factor Rev. 2006;17:305–23.

4. Saltiel AR, Kahn CR. Insulin signalling and the regulation of glucose and lipid metabolism. Nature. 2001;414:799–806.

5. Casa AJ, Dearth RK, Litzenburger BC, Lee A V, Cui X. The type I insulin-like growth factor receptor pathway: a key player in cancer therapeutic resistance. Front Biosci. 2008;13:3273–87.

6. Boone DN, Lee A V. Targeting the insulin-like growth factor receptor: developing biomarkers from gene expression profiling. Crit Rev Oncog. 2012;17:161–73.

7. Renehan AG, Zwahlen M, Egger M. Adiposity and cancer risk: new mechanistic insights from epidemiology. Nat Rev Cancer. 2015;15:484–98.

8. Khandekar MJ, Cohen P, Spiegelman BM. Molecular mechanisms of cancer development in obesity. Nat Rev Cancer. 2011;11:886–95.

9. Pollak MN, Schernhammer ES, Hankinson SE. Insulin-like growth factors and neoplasia. Nat Rev Cancer. 2004;4:505–18.

10. Pollak M. Insulin and insulin-like growth factor signalling in neoplasia. Nat Rev Cancer. 2008;8:915–28.

11. Casa A, Litzenburger B, Dearth R, Lee A v. Insulin-Like Growth Factor Signaling in Normal Mammary Gland Development and Breast Cancer Progression. Breast Cancer Progn Treat Prev. 2008;303.

12. Maki RG. Small is beautiful: insulin-like growth factors and their role in growth, development, and cancer. J Clin Oncol. 2010;28:4985–95.

13. Gallagher EJ, LeRoith D. The proliferating role of insulin and insulin-like growth factors in cancer. Trends Endocrinol Metab. 2010;21:610–8.

14. Siddle K. Signalling by insulin and IGF receptors: supporting acts and new players. J Mol Endocrinol. 2011;47:R1–10.

15. Clayton PE, Banerjee I, Murray PG, Renehan AG. Growth hormone, the insulin-like growth factor axis, insulin and cancer risk. Nat Rev Endocrinol. 2011;7:11–24.

16. Arcaro A. Targeting the insulin-like growth factor-1 receptor in human cancer. Front Pharmacol. 2013;4.

17. Simpson A, Petnga W, Macaulay VM, Weyer-Czernilofsky U, Bogenrieder T. Insulin-Like Growth Factor (IGF) Pathway Targeting in Cancer: Role of the IGF Axis and Opportunities for Future Combination Studies. Target Oncol. 2017;12:571–97.

18. Olmos D, Basu B, de Bono JS. Targeting insulin-like growth factor signaling: rational combination strategies. Mol Cancer Ther. 2010;9:2447–9.

19. DeFronzo RA, Ferrannini E, Groop L, Henry RR, Herman WH, Holst JJ, et al. Type 2 diabetes mellitus. Nat Rev Primer. 2015;1:15019.

20. Sachdev D, Yee D. The IGF system and breast cancer. Endocr Relat Cancer. 2001;8:197–209.

21. Menting JG, Whittaker J, Margetts MB, Whittaker LJ, Kong GKW, Smith BJ, et al. How insulin engages its primary binding site on the insulin receptor. Nature. 2013;493:241–U276.

22. Kavran JM, McCabe JM, Byrne PO, Connacher MK, Wang ZH, Ramek A, et al. How IGF-1 Activates its Receptor. Elife. 2014;3.

23. Houde D, Demarest SJ. Fine Details of IGF-1R Activation, Inhibition, and Asymmetry Determined by Associated Hydrogen/Deuterium-Exchange and Peptide Mass Mapping. Structure. 2011;19:890–900.

24. Kiselyov VV, Versteyhe S, Gauguin L, De Meyts P. Harmonic oscillator model of the insulin and IGF1 receptors’ allosteric binding and activation. Mol Syst Biol. 2009;5.

25. Erdem C, Nagle AM, Casa AJ, Litzenburger BC, Wang Y, Taylor DL, et al. Proteomic Screening and Lasso Regression Reveal Differential Signaling in Insulin and Insulin-like Growth Factor I (IGF1) Pathways. Mol Cell Proteomics. 2016;15:3045–57.

26. Nagle AM, Levine KM, Tasdemir N, Scott JA, Burlbaugh K, Kehm J, et al. Loss of E-cadherin Enhances IGF1–IGF1R Pathway Activation and Sensitizes Breast Cancers to Anti-IGF1R/InsR Inhibitors. Clin Cancer Res. 2018 Oct 15;24(20):5165LP–5177.

27. Bianconi F, Baldelli E, Ludovini V, Crino L, Flacco A, Valigi P. Computational model of EGFR and IGF1R pathways in lung cancer: A Systems Biology approach for Translational Oncology (vol 30, pg 142, 2012). Biotechnol Adv. 2013;31:358–60.

28. Halasz M, Kholodenko BN, Kolch W, Santra T. Integrating network reconstruction with mechanistic modeling to predict cancer therapies. Sci Signal. 2016;9(455):ra114.

29. Sedaghat AR, Sherman A, Quon MJ. A mathematical model of metabolic insulin signaling pathways. Am J Physiol Endocrinol Metab. 2002;283:E1084–101.

30. Borisov N, Aksamitiene E, Kiyatkin A, Legewie S, Berkhout J, Maiwald T, et al. Systems-level interactions between insulin-EGF networks amplify mitogenic signaling. Mol Syst Biol. 2009;5.

31. Bouhaddou M, Barrette AM, Stern AD, Koch RJ, DiStefano MS, Riesel EA, et al. A mechanistic pan-cancer pathway model informed by multi-omics data interprets stochastic cell fate responses to drugs and mitogens. PLoS Comput Biol. 2018;14(3).

32. Tibes R, Qiu Y, Lu Y, Hennessy B, Andreeff M, Mills GB, et al. Reverse phase protein array: validation of a novel proteomic technology and utility for analysis of primary leukemia specimens and hematopoietic stem cells. Mol Cancer Ther. 2006;5:2512–21.

33. Varghese RS, Zuo YM, Zhao Y, Zhang YW, Jablonski SA, Pierobon M, et al. Protein network construction using reverse phase protein array data. Methods. 2017;124:89–99.

34. Harris LA, Hogg JS, Tapia JJ, Sekar JA, Gupta S, Korsunsky I, et al. BioNetGen 2.2: advances in rule-based modeling. Bioinformatics. 2016;32:3366–8.

35. Gupta S, Hainsworth L, Hogg J, Lee R, Faeder J. Evaluation of Parallel Tempering to Accelerate Bayesian Parameter Estimation in Systems Biology. In: 2018 26th Euromicro International Conference on Parallel, Distributed and Network-based Processing (PDP). 2018. p. 690–7.

36. Swigon D. Ensemble Modeling of Biological Systems. In: Antoniouk A V., Melnik RVN, editors. Mathematics and Life Sciences. Walter de Gruyter; 2012. p. 19–42.

37. Brown KS, Sethna JP. Statistical mechanical approaches to models with many poorly known parameters. Phys Rev E. 2003 Aug 12;68(2):21904.

38. Kruger M, Kratchmarova I, Blagoev B, Tseng YH, Kahn CR, Mann M. Dissection of the insulin signaling pathway via quantitative phosphoproteomics. Proc Natl Acad Sci U S A. 2008;105:2451–6.

39. Cai W, Sakaguchi M, Kleinridders A, Gonzalez-Del Pino G, Dreyfuss JM, O’Neill BT, et al. Domain-dependent effects of insulin and IGF-1 receptors on signalling and gene expression. Nat Commun. 2017;8:14892.

40. White MF. The IRS-1 signaling system. Curr Opin Genet Dev. 1994 Feb 1;4(1):47–54.

41. Weng L-P. PTEN inhibits insulin-stimulated MEK/MAPK activation and cell growth by blocking IRS-1 phosphorylation and IRS-1/Grb-2/Sos complex formation in a breast cancer model. Hum Mol Genet. 2002;10(6):605–16.

42. Mendoza MC, Er EE, Blenis J. The Ras-ERK and PI3K-mTOR pathways: cross-talk and compensation. Trends Biochem Sci. 2011 Jun 1;36(6):320–8.

43. Xu W, Smith AM, Faeder JR, Marai GE. RuleBender: a visual interface for rule-based modeling. Bioinformatics. 2011;27:1721–2.

44. Lopez CF, Muhlich JL, Bachman JA, Sorger PK. Programming biological models in Python using PySB. Mol Syst Biol. 2013;9.

45. Danos V, Laneve C. Formal molecular biology. Theor Comput Sci. 2004;325:69–110.

46. Rukhlenko OS, Khorsand F, Krstic A, Rozanc J, Alexopoulos LG, Rauch N, et al. Dissecting RAF Inhibitor Resistance by Structure-based Modeling Reveals Ways to Overcome Oncogenic RAS Signaling. Cell Syst. 2018 Aug;7(2):161–179.e14.

47. Klaeger S, Heinzlmeir S, Wilhelm M, Polzer H, Vick B, Koenig P-A, et al. The target landscape of clinical kinase drugs.

48. Taniguchi CM, Emanuelli B, Kahn CR. Critical nodes in signalling pathways: insights into insulin action. Nat Rev Mol Cell Biol. 2006;7:85–96.

49. Shi T, Niepel M, McDermott JE, Gao Y, Nicora CD, Chrisler WB, et al. Conservation of protein abundance patterns reveals the regulatory architecture of the EGFR-MAPK pathway. Sci Signal. 2016;9:rs6.

50. Stern AM, Schurdak ME, Bahar I, Berg JM, Taylor DL. A Perspective on Implementing a Quantitative Systems Pharmacology Platform for Drug Discovery and the Advancement of Personalized Medicine. J Biomol Screen. 2016;21:521–34.

51. Musante CJ, Ramanujan S, Schmidt BJ, Ghobrial OG, Lu J, Heatherington AC. Quantitative Systems Pharmacology: A Case for Disease Models. Clin Pharmacol Ther. 2017;101:24–7.

52. Zhao S, Iyengar R. Systems pharmacology: network analysis to identify multiscale mechanisms of drug action. Annu Rev Pharmacol Toxicol. 2012;52:505–21.

